# The neural control of infanticide and parental behaviors in male mice

**DOI:** 10.64898/2026.03.12.711470

**Authors:** Long Mei, Yifan Wang, Qingfeng Wei, Yujie Xiong, Patrick T. O’Neill, Bingxiang Yang, Dayu Lin

## Abstract

Infanticide, the killing of conspecific young, is a natural behavior observed commonly in non-parental animals across species, including mice. Our recent study in female mice revealed a mutually inhibitory circuit, composed of estrogen receptor alpha cells in the medial preoptic area (MPOA^Esr1^) and the posterior part of the bed nucleus of stria terminalis (BNSTp^Esr1^), that controls pup-directed behaviors, with the former driving maternal care and the latter promoting infanticide. Given that both MPOA and BNSTp are sexually dimorphic, here we asked whether the same circuit operates in males. Our functional manipulations and in vivo and in vitro recordings reveal that MPOA^Esr1^ and BNSTp^Esr1^ cells similarly and respectively drive paternal care and infanticide and antagonize each other in male mice. Furthermore, during fatherhood, MPOA^Esr1^ cell excitability increases while BNSTp^Esr1^ cell excitability decreases to enable the switch from infanticide to paternal care. Despite the similarity in circuit organization, a direct comparison between males and females reveals sex differences in the intrinsic properties of MPOA^Esr1^ and BNSTp^Esr1^ cells. Thus, MPOA^Esr1^-BNSTp^Esr1^ emerges as a common circuit motif for controlling pup-directed behaviors in both sexes, whereas the activity balance between these two populations differs between sexes and likely contributes to the different tendencies in males and females to express pup-caring versus killing behaviors.

## Introduction

Parental behavior is essential for the survival and well-being of offspring, but demands significant energy expenditure^1^. To optimize the use of limited energy, parental behavior is typically heightened only during specific reproductive phases, such as when an individual becomes a father or mother^2–6^. Outside these periods, individuals often exhibit neutral or even aggressive behavior toward the young^2–6^. The transition from a low to a high pup-caring state during parenthood is crucial for the survival of the young.

Extensive research has identified key brain regions supporting pup-directed behaviors. The medial preoptic area (MPOA) was identified as a key regulator of paternal behavior since 1970^7–10^. In the last decade, the neural substrates underlying infanticidal behavior have become a growing field of research^11–16^. Several regions and cell populations were found effective in modulating infanticide in male mice, including the rhomboid nucleus of the bed nucleus of stria terminalis (BNSTrh)^13^, GABAergic cells in the medial amygdala posterodorsal part (MeApd)^16^, urocortin-3 positive cells in the perifornical area (PFA^Ucn3^)^12^, and the amygdalohippocampal area (also known as posterior amygdala) cells projecting to the medial preoptic area (AHi^→MPOA^)^15^. Most recently, we identified estrogen receptor alpha-expressing (Esr1) cells in the posterior part of the BNST (BNSTp) as a key population to drive infanticide in female mice^14^. BNSTp^Esr1^ cells are highly active during infanticide but not maternal care and can bi-directionally control acute pup-directed attack^14^. Furthermore, BNSTp^Esr1^ and MPOA^Esr1^ cells are antagonistic to each other through direct inhibitory synaptic connection^9,14^.

Notably, the results so far suggest that the neural substrates for infanticide have little in common between the sexes. In males, BNSTrh, MEApd^Vgat^, PFA^Ucn3^, and AHi ^(MPOA^ ^cells)^ are known to modulate infanticide, while in females, BNSTp^Esr1^ is the key player. Is the infanticide circuit sex-specific? A positive answer appears to be reasonable, given that many identified infanticide-relevant regions are sexually dimorphic. For example, the projection pattern of AHi in the MPOA differs substantially between sexes: female, but not male, AHi targets AVPV densely^17^. The male BNSTp is nearly twice the size of the female BNSTp, although Esr1 is more abundantly expressed in females than males^14,18,19^. However, one caveat of this conclusion is that almost all recent studies on infanticide focused on one sex, leaving the function of the cells in the other sex unexplored. Thus, the seemingly sex-specific infanticide circuit could be simply a result of experimental subject bias.

To address whether the infanticide circuit is distinct between sexes, we followed up on our recent study in females^14^ and examined the function of BNSTp^Esr1^ cells in male infanticide and their relationship with MPOA^Esr1^ cells. Additionally, we compared the cellular properties of BNSTp^Esr1^ and MPOA^Esr1^ cells between females and males, as well as between virgins and parents.

## Results

### Responses of male BNSTp^Esr1^ cells during infanticide and parental behaviors

In our recent study^14^, we reported that infanticidal behaviors in female mice differed drastically across strains. While nearly no C57BL/6 (C57) virgin female mice (2/165) spontaneously attacked pups, approximately one-third (50/146) virgin Swiss Webster (SW) female mice attacked and killed pups^14^. In males, the strain difference is less prominent. Both C57 and SW virgin male mice reportedly show markedly higher rates of pup-killing than females^20,21^. Here, we found that 40% (54/134) of C57 and 71% (58/82) of SW virgin males attacked pups during the 10-minute test session. During fatherhood, no male, regardless of their genetic background, attacked pups. Instead, almost all males showed parental behaviors, including pup retrieval, crouching, and grooming pups. In this study, we mainly used SW male mice as our test subjects to be consistent with our previous study in females^14^. C57 mice were also used in some cases to confirm the generality of our findings across strains.

To understand the responses of BNSTp^Esr1^ cells in male infanticide, we recorded the Ca^2+^ activity of BNSTp^Esr1^ cells in sexually naïve male and fathering mice using fiber photometry. Specifically, we virally expressed GCaMP6f^22^, a genetically encoded Ca^2+^ sensor, in BNSTp^Esr1^ cells of Esr1-2A-Cre virgin SW male mice and implanted a 400-μm optic fiber above the injection site (**Fig. 1a-c**). Before surgery, animals were screened for infanticide, and only the ones that spontaneously attacked pups were used. Three weeks after surgery, animals were recorded during pup interaction. Then, each male was paired with a female mouse and became a father. Three to four days after female parturition, BNSTp^Esr1^ activities were recorded again when the males expressed paternal behaviors (**Fig. 1d**).

**Figure 1.**
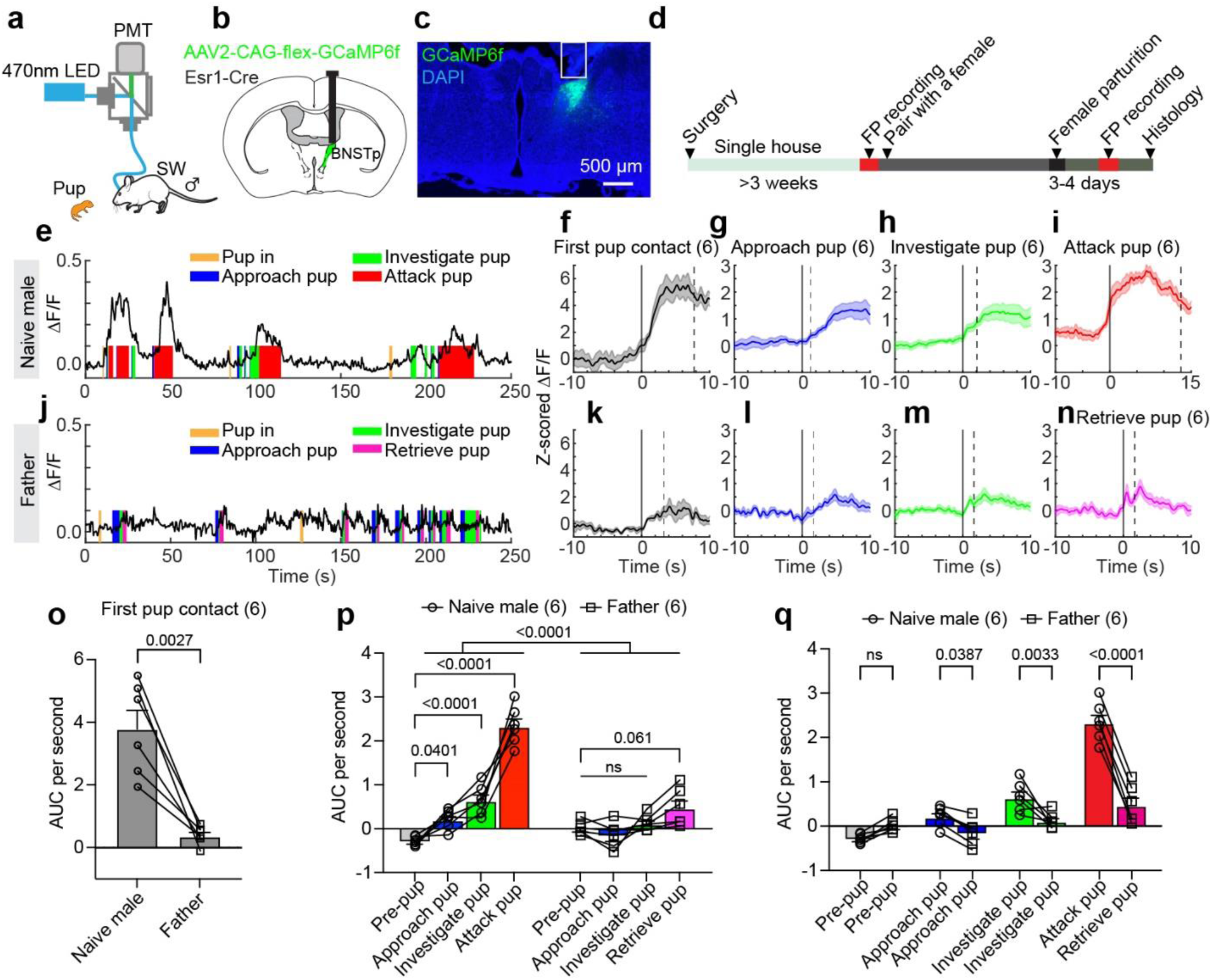
The response pattern of BNSTp^Esr1^ cells during male infanticide and paternal care. **(a)** The fiber photometry setup. **(b)** The illustration of targeting BSNTp^Esr1^ cells for fiber photometry recording. **(c)** A representative image showing the fiber track (white line) and GCaMP6f (green) expression in the BSNTp. **(d)** The experimental timeline. **(e and j)** Representative ΔF/F traces of BNSTp^Esr1^ cells during pup interaction of a hostile naïve male (**e**) and a father (**j**). **(f-i)** PETHs of Z-scored ΔF/F of BNSTp^Esr1^ cells aligned to the onset of first pup contact (**f**), pup approach (**g**), pup investigation (**h**), and pup attack (**i**) of hostile naïve males. (n=6 mice, data shown as mean ± SEM.) **(k-n)** PETHs of Z-scored ΔF/F of BNSTp^Esr1^ cells aligned to the onset of first pup contact (**k**), approach pup (**l**), investigate pup (**m**), and retrieve pup (**n**) of fathers. (n=6 mice, data shown as mean ± SEM.) **(o)** The average area under the curve (AUC) of Z-scored ΔF/F per second during the first pup contact. (n=6 male mice. Paired t-test; p=0.0027. Data shown as mean + SEM.) **(p)** The AUC of the Z-scored ΔF/F per second during various pup-directed behaviors to compare responses across behaviors in hostile naïve males and fathers. (n=6 mice. Two-way RM ANOVA followed by Bonferroni’s multiple comparisons test; Naive-Father, p<0.0001; Naive male: Pre-pup vs. Approach pup, p=0.0401; Pre-pup vs. Investigate pup, p<0.0001; Pre-pup vs. Attack pup, p<0.0001; Father: Pre-pup vs. Retrieve pup, p=0.061; ns: not significant, data shown as mean + SEM.) **(q)** The AUC of the Z-scored ΔF/F per second during each pup-directed behavior to compare responses between hostile naïve males and fathers. (n=6 mice. Two-way RM ANOVA followed by Bonferroni’s multiple comparison tests; Pre-pup: Naïve male vs. Father, ns: not significant; Approach pup: Naïve male vs. Father, p=0.0387; Investigate pup: Naïve male vs. Father, p=0.0033; Attack pup: Naïve male vs. Father, p<0.0001. Data shown as mean + SEM.)

In naïve males, GCaMP signals increased sharply during the first pup contact (**Fig. 1e-f**). The signal also increased during the subsequent pup approach and investigation and peaked during pup attack (**Fig. 1g-i)**. In contrast, we observed minimal GCaMP signal increase during pup introduction in fathers or when the males approached, investigated, and retrieved pups (**Fig. 1j-n**). Overall, the average GCaMP signal significantly increased from the pre-pup baseline during pup approach, investigation, and attack in naïve males, while there was no significant signal increase during pup-directed behaviors in fathers (**Fig. 1p**). When comparing different reproductive states, the neural response was significantly higher in infanticidal naïve males than in fathers during first pup contact and various pup-directed behaviors (**Fig. 1o, q**). In comparison to the impressive activity increase of BNSTp^Esr1^ cells during pup attack, attacking an adult male evoked only a slight increase in GCaMP signal (**Supplementary Fig. 1**). These results suggest that BNSTp^Esr1^ cells are more active during hostile infant-directed behaviors than positive pup-directed behaviors in males, as is the case in females^14^.

### BNSTp^Esr1^ plays an indispensable role in male infanticide behavior

To understand whether BNSTp^Esr1^ cells are functionally important in driving male infanticide, we expressed diphtheria toxin receptor (DTR) in BNSTp^Esr1^ neurons bilaterally of naïve Esr1-2A-Cre SW male mice (**Fig. 2a**). Control animals expressed mCherry in BNSTp^Esr1^ neurons (**Fig. 2a**). All males were screened before the surgery. Only animals that showed spontaneous infanticide were used. Three weeks after surgery, we again tested their behavior towards pups and confirmed that all males remained infanticidal (**Fig. 2b**). We then injected 50ug/kg diphtheria toxin (DT) intraperitoneally to induce apoptosis of DTR-expressing cells^23^. Histology analysis confirmed successful ablation of Esr1-positive cells in the BNSTp but not the more ventrally situated MPOA in DTR animals (**Fig. 2c, d**). One week after the DT injection, we again tested the pup-directed behaviors by introducing 1-3 P1-P4 pups into the testing mouse cage. While 8/8 mCherry control mice continued to attack pups quickly, only 2/8 DTR mice did so, and 5/8 DTR mice even retrieved pups (**Fig. 2e-j**). The latency to investigate pups did not change after DT injection (**Fig. 2k**).

**Figure 2.**
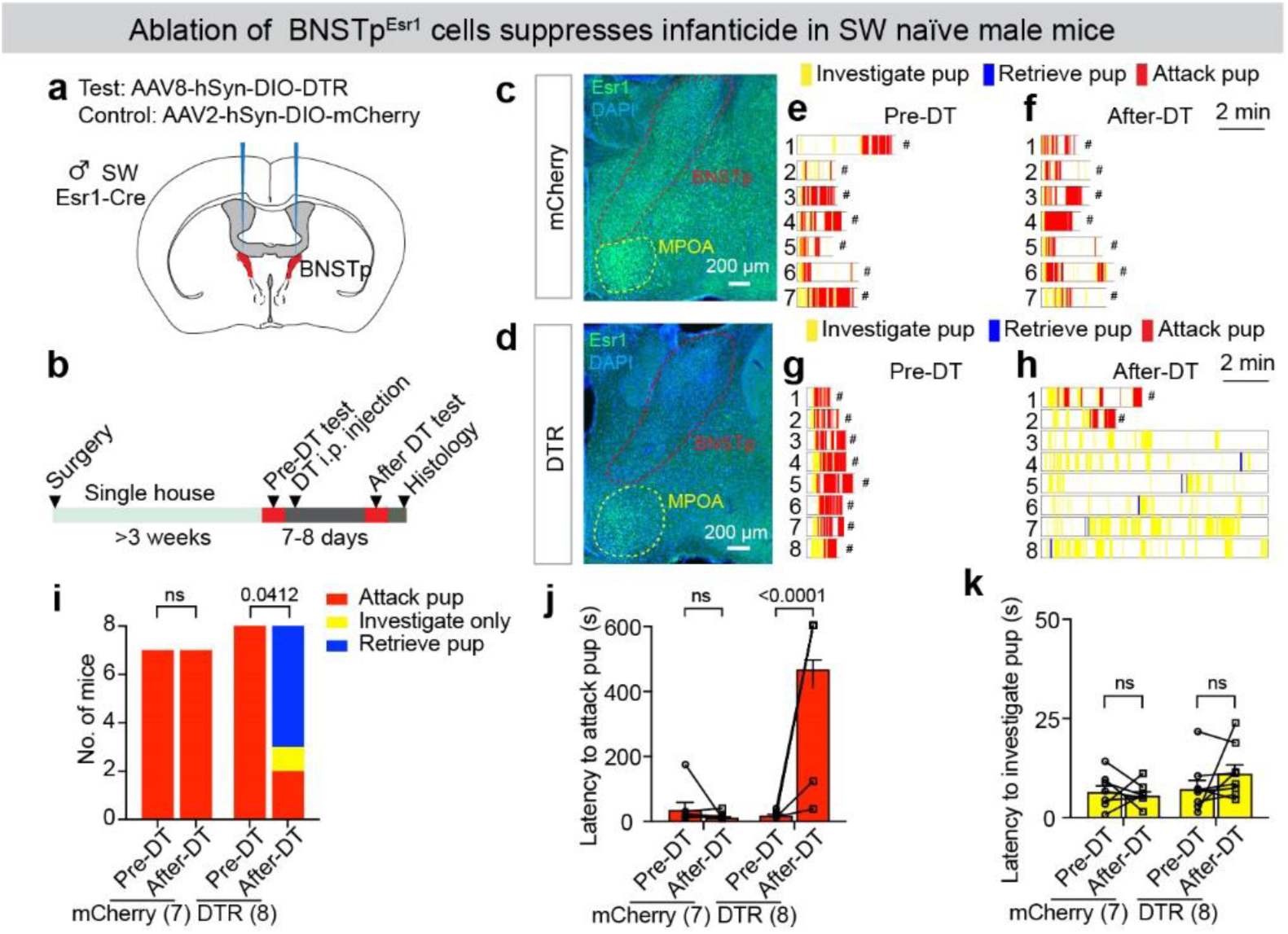
BNSTp^Esr1^ neurons are necessary for male infanticide. **(a)** The strategy to ablate BNSTp^Esr1^ cells in SW Esr1-Cre males. **(b)** The experimental timeline. **(c and d)** Representative images showing Esr1 staining (green) in mCherry (**c**) and DTR (**d**) expressing mice. **(e and f)** Raster plots showing pup-directed behaviors in mCherry male mice before **(e)** and after (**f**) DT injection. # wounded pups were removed and euthanized. **(g and h)** Raster plots showing pup-directed behaviors in DTR male mice before (g) and after (h) DT injection. # wounded pups were removed and euthanized. **(i)** The number of mCherry and DTR male mice that attacked, investigated only, or retrieved pups before and after DT injection. (McNemar’s test; ns: not significant; p=0.0412.) **(j)** Latency to attack pup of mCherry and DTR male mice before and after DT injection. The latency equals 600 s if no pup attack occurs during the 10-minute test. (n = 7 males of mCherry group, n = 8 males of DTR group. Mixed effect two-way ANOVA followed by Bonferroni’s multiple comparisons test; ns: not significant; p <0.0001. Data shown as mean + SEM.) **(k)** Latency to investigate pup of mCherry and DTR male mice before and after DT injection. (n = 7 males of mCherry group; n = 8 males of DTR group. Mixed effect two-way ANOVA followed by Bonferroni’s multiple comparisons test, ns: not significant. Data shown as mean + SEM.)

We also confirmed the necessity of BNSTp^Esr1^ cells in C57BL/6 naïve male mice using chemogenetic inhibition (**Supplementary Fig. 2a, b**). After saline injection, both hM4Di and mCherry control male mice attacked pups quickly (**Supplementary Fig. 2c-f**). After CNO injection, only 1/7 hM4Di male mice attacked the pup after a long delay, while all mCherry animals continued to show infanticide (**Supplementary Fig. 2c-f**). The decreased pup attack was not due to a lack of pup interaction, as the latency to pup investigation did not differ between saline and CNO-injected days (**Supplementary Fig. 2g**).

Furthermore, we found that inhibition of BNSTp^Esr1^ cells suppressed male aggression, as previously reported^24^ (**Supplementary Fig. 2h-m**). Although all test males attacked the adult male intruder after both saline and CNO injection, the attack latency was significantly longer, and the attack duration decreased by approximately 50% after CNO injection compared to saline days (**Supplementary Fig. 2k-m**). These data support the indispensable role of BNSTp^Esr1^ cells in driving male infanticide and a minor but still important role in promoting inter-male aggression.

### BNSTp^Esr1^ neurons are sufficient to induce infanticide in male mice

To understand whether BNSTp^Esr1^ cells are sufficient to promote male infanticide, we virally expressed Channelrhodopsin-2 (ChR2) in bilateral BNSTp^Esr1^ neurons in naïve SW male mice and optogenetically activated the cells during pup interaction (**Fig. 3a-c**). Control animals were injected with GFP viruses (**Fig. 3a**). All males were screened before and after the surgery, and only animals that did not show spontaneous infanticide were used. During the test, we placed 1-2 pups in the test male’s home cage and delivered sham (0 mW) or blue light (0.5-2 mW, 20ms, 20Hz) for 20s when the male investigated a pup (**Fig. 3b**). We found that optogenetic activation of BNSTp^Esr1^ neurons induced pup attack robustly in all ChR2 males, while none of the GFP control males attacked the pup during the recording session (**Fig. 3d, h**). In 90% of light trials but none of the sham trials, the ChR2 males attacked pups, with an average latency of less than 2 seconds (**Fig. 3e-f, h-j**).

**Figure 3.**
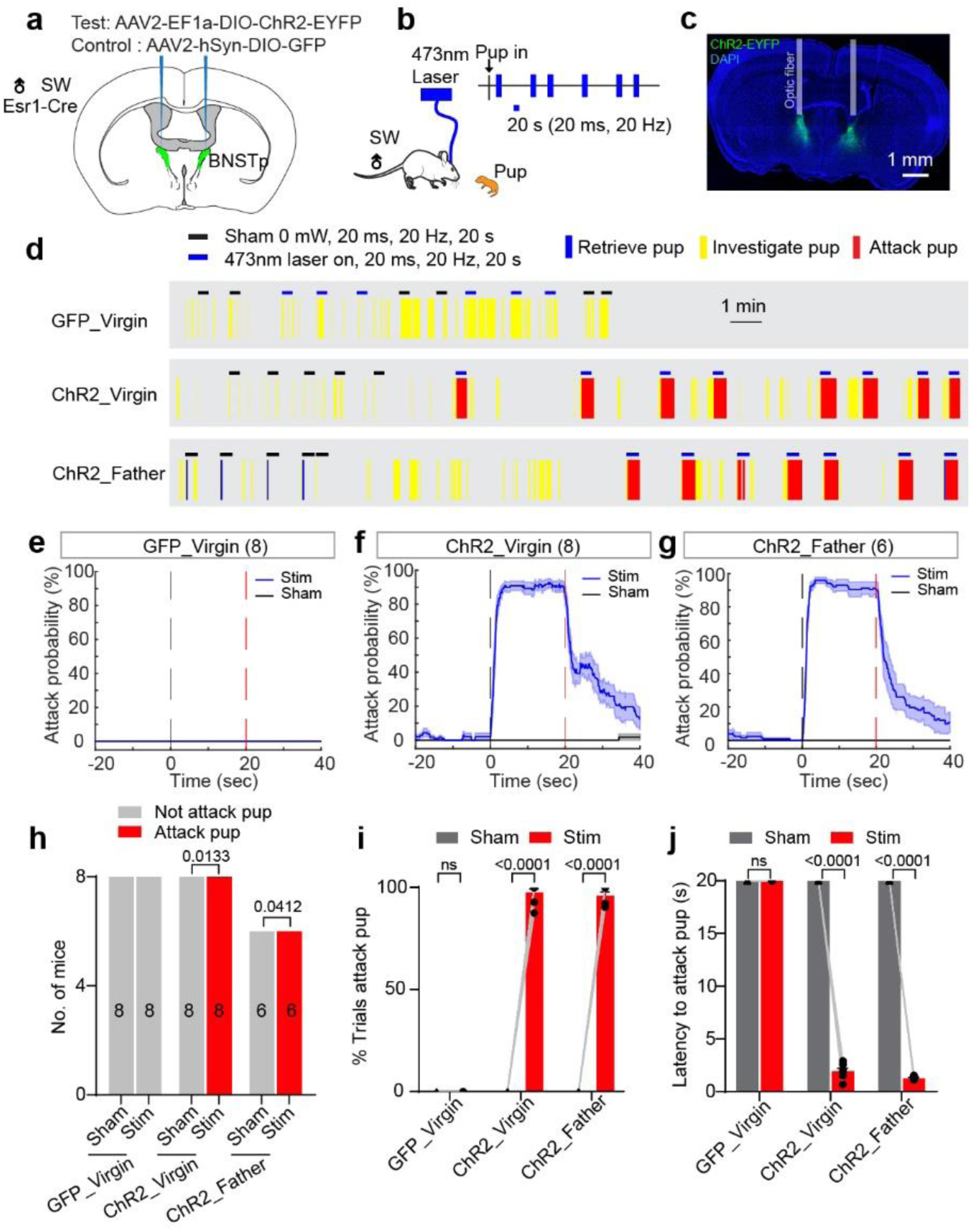
Activating BNSTp^Esr1^ cells is sufficient to induce infanticide in males. **(a and b)** The strategy to optogenetically activate BNSTp^Esr1^ cells. **(c)** A representative image showing ChR2 (green) expression in the BNSTp and optic fiber locations. **(d)** Raster plots showing pup-directed behaviors during sham and light stimulation. **(e-g)** PETHs of attacking pup probability aligned to sham (black) and light (blue) onset of GFP virgin male (**e**), ChR2 virgin male (**f**), and ChR2 father males (**g**). The dashed lines mark the trial period. (n=8 males for GFP virgin and ChR2 virgin group, n=6 males for ChR2 father group. Data shown as mean ± SEM.) **(h)** Number of males that attacked pups during the stimulation periods. (n=8 males for GFP virgin and ChR2 virgin group, n=6 males for ChR2 father group. McNemar’s test.) **(i)** The percentage of trials in which the male attacked pups. (n=8 males for GFP virgin and ChR2 virgin group, n=6 males for ChR2 father group. Mixed effect two-way ANOVA followed by Bonferroni’s multiple comparisons test, ns: not significant. Data shown as mean + SEM) **(j)** The average latency to attack pup after sham or light stimulation onset. The latency equals 20 s if no attack occurs during the 20s sham or light stimulation period. (n=8 males for GFP virgin and ChR2 virgin group, n=6 males for ChR2 father group. Mixed effect two-way ANOVA followed by Bonferroni’s multiple comparisons test, ns: not significant. Data shown as mean + SEM.)

To determine whether the induced infanticidal behavior is specific to the naïve state, we paired each ChR2 male with a female. 6/8 paired females successfully gave birth. Two to four days after the test males became fathers, we optogenetically activated BNSTp^Esr1^ neurons during pup interactions and observed time-locked pup attack (**Fig. 3d, g**). All 6 tested fathering males attacked pups rapidly (Latency to attack < 2 seconds) in over 90% of light-on trials, whereas no attack ever occurred during sham (no light) trials (**Fig. 3h–j**).

BNSTp^Esr1^ cells in males have been indicated in inter-male aggression^25^. To address whether infanticide induced by BNSTp^Esr1^ neuron activation represents target-unspecific attacks, we optogenetically activated BNSTp^Esr1^ cells in the presence of a non-aggressive adult Balb/C male intruder and found no light-induced attack (**Supplementary Fig. 3a, b, d**). Instead, the stimulated animal spent significantly more time grooming the Balb/C males (**Supplementary Fig. 3a, c, e**). Of note, while the previous study showed inhibiting BNSTp^Esr1^ cells can disrupt ongoing inter-male aggression, stimulation-locked adult-directed attack has never been reported^25^.

As a complementary strategy, we chemogenetically activated BNSTp^Esr1^ neurons by bilaterally expressing hM3Dq-mCherry in BNSTp^Esr1^ neurons in naïve Esr1-2A-Cre non-infanticidal SW male mice and found that all test animals attacked the pups after CNO but not saline injections (**Supplementary Fig. 4**).

These results suggest that BNSTp^Esr1^ neurons are highly effective in driving infanticidal behavior in male mice and that the induced aggression is directed specifically toward pups rather than reflecting a general increase in aggressive behavior.

### MPOA^Esr1^ is naturally activated during paternal behavior but not infanticide in male mice

MPOA^Esr1^ cells are essential for driving parental behaviors in female mice, and they antagonize BNSTp^Esr1^ cell activity to suppress female infanticide^14^. We next asked whether the same BNSTp^Esr1^-MPOA^Esr1^ circuit exists in males. To begin, we examined the responses of MPOA^Esr1^ cells in parental behaviors in males. Of note, multiple functional studies have suggested that MPOA is similarly important for parental behaviors in males as in females (but see Dimén et al.^26^). For example, optogenetic activation of MPOA^Esr1^ cells can induce pup retrieval in both male and female mice^9,27^. Ablating MPOA galanin cells (a population partially overlapping with Esr1 cells) significantly impairs pup retrieval and other parental behaviors in fathers^8^. Deleting prolactin receptors in the MPOA of male mice impairs all aspects of parental behaviors in fathering mice^28^. Nevertheless, none of these recent studies examined the in vivo responses of MPOA cells to pups as the animals switch from infanticide to paternal care during fatherhood.

We used fiber photometry to record population Ca^2+^ activity of MPOA^Esr1^ cells in Esr1-2A-Cre SW male mice (**Fig. 4a-c**). After recording the Ca^2+^ response in naïve infanticidal males, we paired each male with a female and recorded again three to four days after they became a father (**Fig. 4d**). In naïve infanticidal males, Ca^2+^ signal started to rise when the male approached the pup (**Fig. 4e, f**), reaching a peak about one second after investigating the pup, and then declined during the later phase of pup investigation (**Fig. 4g**). During subsequent pup attacks, Ca^2+^ signal was mildly suppressed (**Fig. 4h**). In contrast, when the males became fathers and exhibited paternal behavior toward pups, Ca^2+^ signal began to rise as they approached the pup (**Fig. 4i, j**), continued increasing throughout pup investigation (**Fig. 4k**), peaked at the onset of pup retrieval, and remained at a high level throughout the retrieval period (**Fig. 4l**). Overall, MPOA^Esr1^ cells in males showed higher activity during paternal behaviors and a trend of suppression during pup attacks (**Fig. 4m, n**). This activity pattern contrasts with that of BNSTp^Esr1^ cell responses to pups (**Fig. 1e-q**).

**Figure 4.**
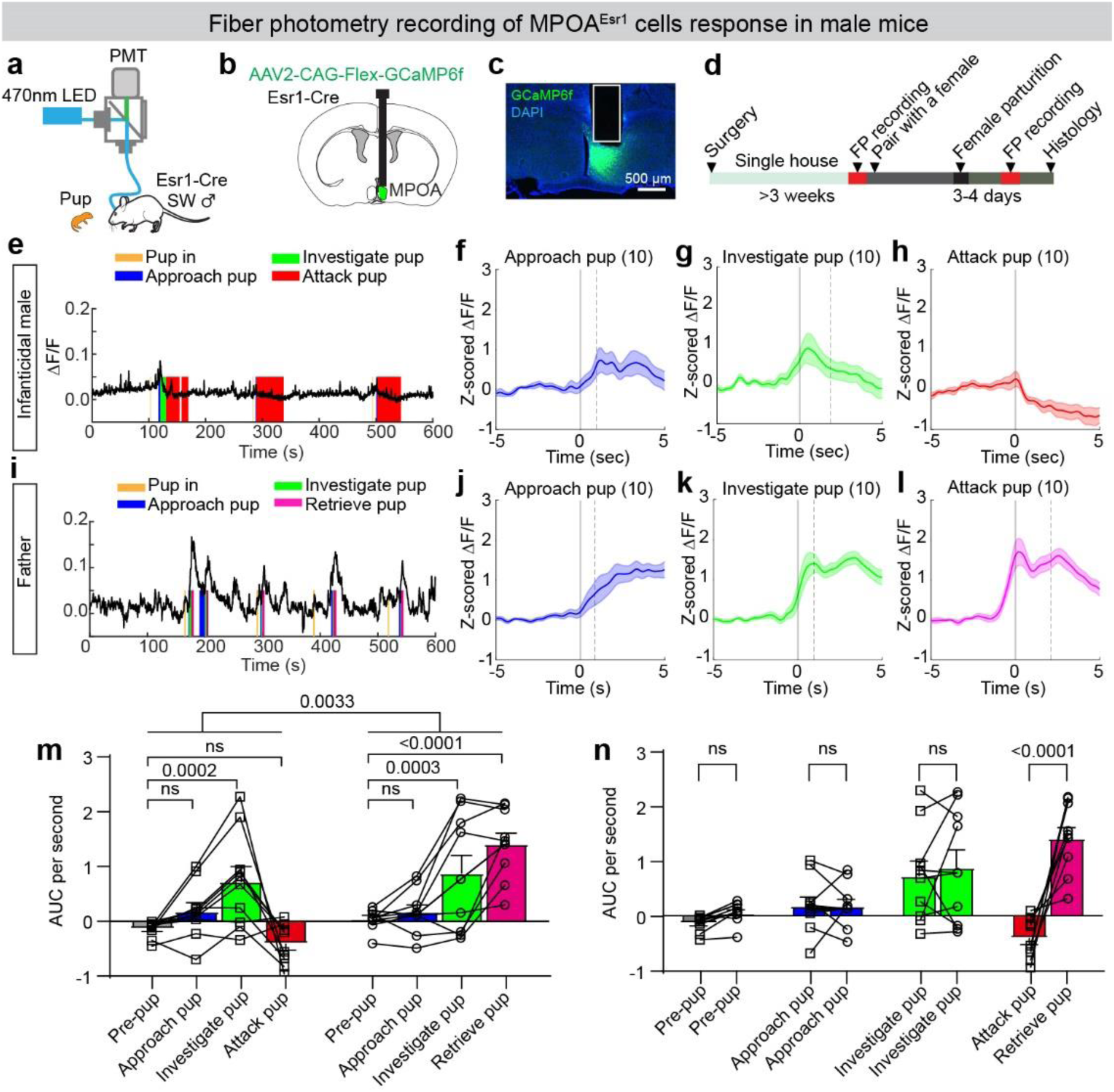
The response pattern of MPOA^Esr1^ cells during male infanticide and paternal care. **(a)** The fiber photometry setup. **(b)** The illustration of targeting MPOA^Esr1^ cells for fiber photometry recording. **(c)** A representative image showing the fiber track (white line) and GCaMP6f (green) expression in the MPOA. **(d)** The experimental timeline. **(e and i)** Representative ΔF/F traces of MPOA^Esr1^ cells during pup interaction of a hostile naïve male (**e**) and a father (**i**). **(f-h)** PETHs of Z-scored ΔF/F of MPOA^Esr1^ cells aligned to the onset of pup approach **(f)**, pup investigation (**g**), and pup attack (**h**) of hostile naïve males. (n=10 mice, data shown as mean ± SEM.) **(j-l)** PETHs of Z-scored ΔF/F of MPOA^Esr1^ cells aligned to the onset of approach pup (**j**), investigate pup (**k**), and retrieve pup (**l**) of fathers. (n=10 mice, data shown as mean ± SEM.) **(m)** The AUC of the Z-scored ΔF/F per second during various pup-directed behaviors to compare responses across behaviors in hostile naïve males and fathers. (n=10 mice. Two-way RM ANOVA followed by Bonferroni’s multiple comparisons test; ns: not significant, data shown as mean + SEM.) **(n)** The AUC of the Z-scored ΔF/F per second during each pup-directed behavior to compare responses between hostile naïve males and fathers. (n=10 mice. Two-way RM ANOVA followed by Bonferroni’s multiple comparison tests; ns: not significant; data shown as mean + SEM.)

### MPOA^Esr1^ cells negatively modulate infanticide in male mice

If the MPOA^Esr1^ cells antagonize BNSTp^Esr1^ cells, we expect that manipulating MPOA^Esr1^ cell activity will have behavioral effects opposite to those of BNSTp^Esr1^ cells. Consistent with this hypothesis, we found that chemogenetic inhibition of MPOA^Esr1^ cells in naïve non-infanticidal SW males significantly increased the probability of infanticide (**Fig. 5a-e**). Specifically, after saline injection, none of the 9 test mice attacked pups, whereas 6/9 attacked pups after CNO injection (**Fig. 5c-e**). The latency to investigate pups did not differ significantly between saline- and CNO-injected days (**Fig. 5f**).

**Figure 5.**
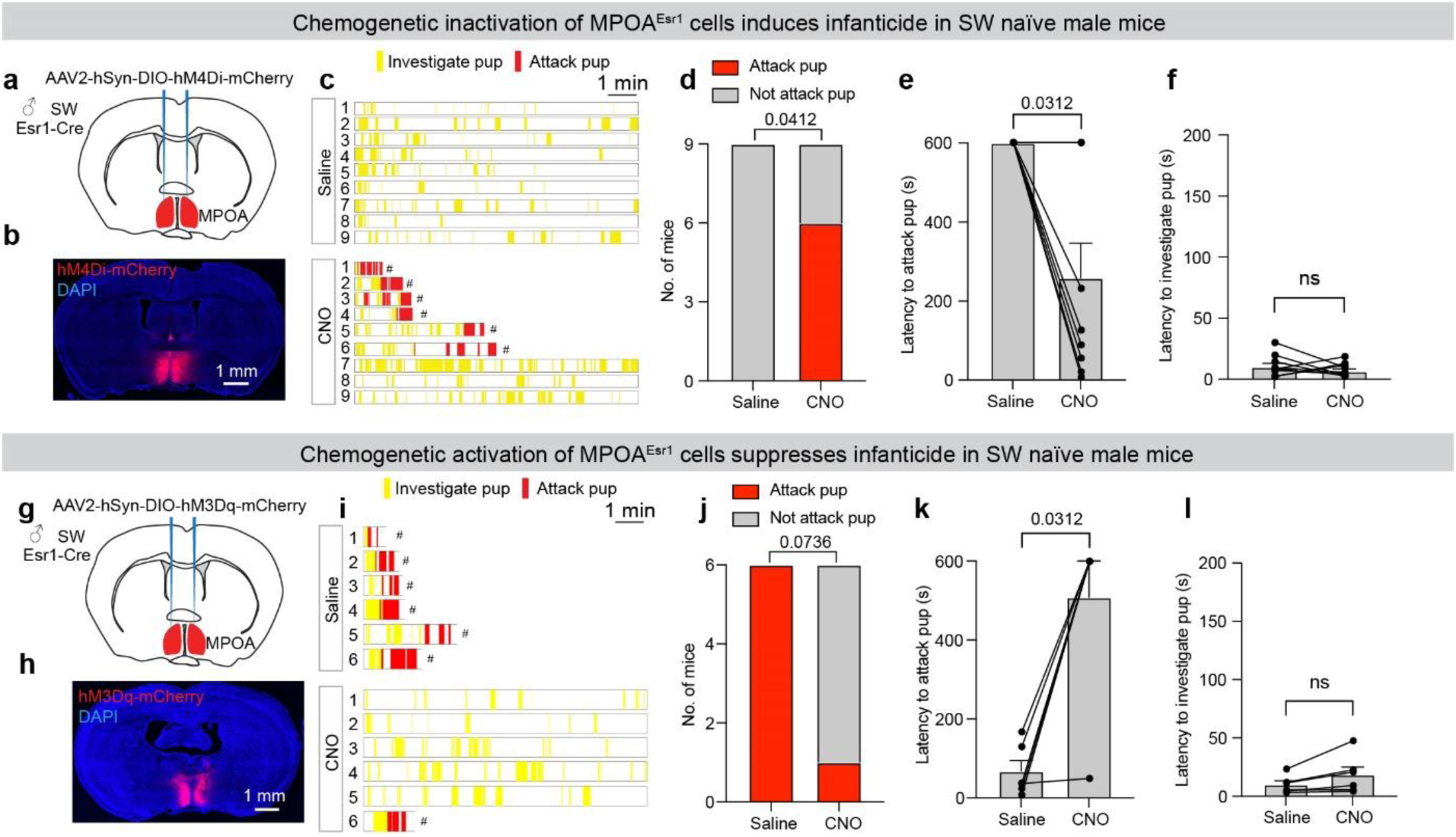
MPOA^Esr1^ neurons bidirectionally regulate infanticidal behavior in sexually naïve male mice. **(a)** The strategy to target MPOA^Esr1^ cells for chemogenetic inhibition. **(b)** A representative image showing hM4Di (red) expression in the bilateral MPOA. **(c)** Raster plots showing pup-directed behaviors after saline or CNO injection. # wounded pups were removed and euthanized. **(d)** The number of males that attacked the pups after saline or CNO injection. (McNemar’s test, p=0.0412.) **(e)** The latency to attack pups after saline or CNO injection. The latency equals 600s if no attack occurs during the 10-minute testing period. (n=9 males. Wilcoxon test; p=0.0312. Data shown as mean + SEM.) **(f)** The latency to investigate the pup after saline or CNO injection. (n=9 males. Paired t-test; ns: not significant. Data shown as mean + SEM.) **(g)** The strategy to target MPOA^Esr1^ cells for chemogenetic activation. **(h)** A representative image showing hM3Dq (red) expression in the bilateral MPOA. **(i)** Raster plots showing pup-directed behaviors after saline or CNO injection. # wounded pups were removed and euthanized. **(j)** The number of males that attacked the pup after saline or CNO injection. (McNemar’s test, p=0.0736.) **(k)** The latency to attack pups after saline or CNO injection. The latency equals 600s if no attack occurs during the 10-minute testing period. (n=6 males. Wilcoxon test; p=0.0312. Data shown as mean + SEM.) **(l)** The latency to investigate the pup after saline or CNO injection. (n=6 males. Paired t-test; ns: not significant. Data shown as mean + SEM.)

Conversely, when we chemogenetically activated MPOA^Esr1^ cells in naïve infanticidal SW males, infanticide was significantly decreased (**Fig. 5g-k**). While all 6 male mice attacked pups after saline injection, only one did so after CNO injection (**Fig. 5i-k**). The pup investigation latency remained short and unchanged after CNO injection (**Fig. 5l**). These results suggest that BNSTp^Esr1^ and MPOA^Esr1^ cells modulate infanticide in opposite directions: while the former promotes the behavior, the latter suppresses it.

### Opposite changes of MPOA^Esr1^ and BNSTp^Esr1^ cell excitability during fatherhood

MPOA^Esr1^ cells increase response to pups during fatherhood, while BNSTp^Esr1^ cells decrease their pup responses (**Fig. 1p and 4m**). To elucidate the physiological mechanisms potentially responsible for the *in vivo* pup response changes during fatherhood, we performed *in vitro* current-clamp recordings from MPOA^Esr1^ and BNSTp^Esr1^ cells in naïve infanticidal males and parental fathers (within 3 days of their partners’ parturition) (**Fig. 6a-f**). For each animal, recordings were obtained from both MPOA^Esr1^ and BNSTp^Esr1^ cells to control for individual variability.

**Figure 6.**
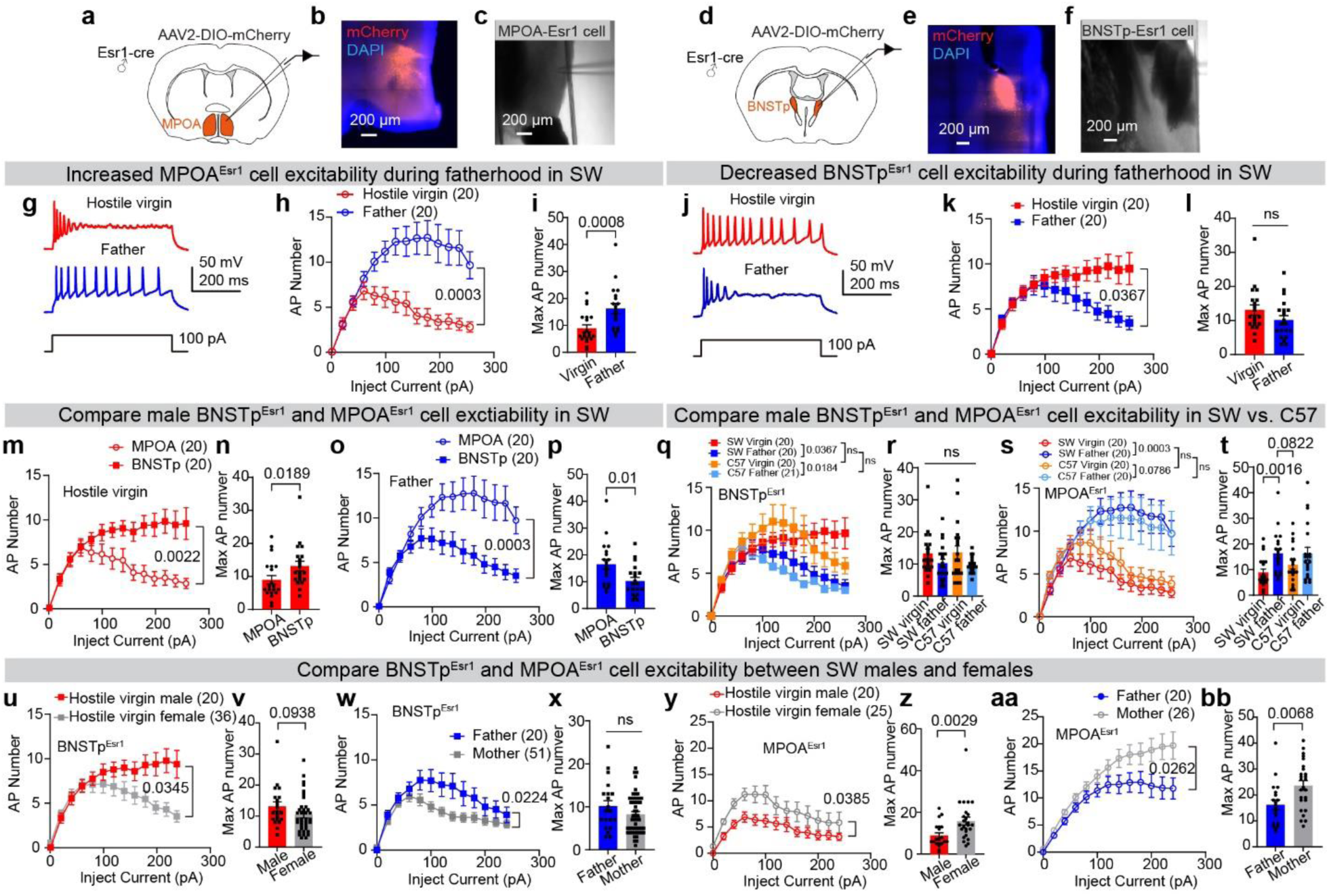
MPOA^Esr1^ and BNSTp^Esr1^ cell excitability change with reproductive states and differ between sexes. **(a and d)** Schematic of in vitro patch-clamp recording from MPOA^Esr1^ cells (**a**) and BNSTp^Esr1^ cells (**d**). **(b and e)** Representative images showing mCherry expression in MPOA^Esr1^ (**b**) and BNSTp^Esr1^ cells (**e**). **(c and f)** Representative images showing the locations of the recorded MPOA^Esr1^ cell and BNSTp^Esr1^ cell (**f**). **(g)** Representative recording traces of MPOA^Esr1^ cells in SW hostile males and fathers. **(h)** *F–I* curves of MPOA^Esr1^ cells from SW hostile virgin males (red, 20 cells from 4 male mice) and fathers (blue, 20 cells from 4 male mice). Data shown as mean ± SEM. Statistical analysis was performed using Two-way RM ANOVA followed by Bonferroni’s multiple comparisons test. **(i)** The maximum number of action potentials of MPOA^Esr1^ cells as shown in (**h**). Data shown as mean ± SEM, two-tailed Mann-Whitney test. **(j)** Representative recording traces of BNSTp^Esr1^ cells in SW hostile virgin males and fathers. **(k)** *F–I* curves of BNSTp^Esr1^ cells from SW hostile virgin males (red, 20 cells from 4 male mice) and fathers (blue, 20 cells from 4 male mice). Statistical analysis was performed using Two-way RM ANOVA followed by Bonferroni’s multiple comparisons test. **(l)** The maximum action potential number of BNSTp^Esr1^ cells as shown in (**k**). Data shown as mean ± SEM, two-tailed unpaired t-test. **(m)** *F–I* curves of MPOA^Esr1^ cells (20 cells from 4 male mice) and BNSTp^Esr1^ cells (20 cells from 4 male mice) from hostile virgin males. Data shown as mean ± SEM. Statistical analysis was performed using Two-way RM ANOVA followed by Bonferroni’s multiple comparisons test. **(n)** The maximum action potential number of MPOA^Esr1^ cells as shown in (**m**). Data shown as mean ± SEM. Two-tailed Mann-Whitney test. **(o)** *F–I* curves of MPOA^Esr1^ cells (20 cells from 4 male mice) and BNSTp^Esr1^ cells (20 cells from 4 male mice) from fathers. Data shown as mean ± SEM. Two-way RM ANOVA followed by Bonferroni’s multiple comparisons test. **(p)** The maximum action potential number of MPOA^Esr1^ cells as shown in (**o**). Data shown as mean ± SEM. Two-tailed Unpaired t-test. **(q)** *F–I* curves of BNSTp^Esr1^ cells from SW and C57 hostile virgin males (SW: red, 20 cells from 4 male mice; C57: yellow, 20 cells from 3 male mice) and fathers (SW: blue, 20 cells from 4 male mice; C57: Cyan, 21 cells from 4 male mice). Data shown as mean ± SEM. Statistical analysis was performed using two-way RM ANOVA followed by Bonferroni’s multiple comparisons test. **(r)** The maximum action potential number of the BNSTp^Esr1^ cells as shown in **(q)**. Data shown as mean ± SEM. Two-tailed Unpaired Ordinary one-way ANOVA followed by Bonferroni’s multiple comparisons test. **(s)** *F–I* curves of MPOA^Esr1^ cells from SW and C57BL/6 hostile virgin males (SW: red, 20 cells from 4 male mice; C57: yellow, 20 cells from 3 male mice) and fathers (SW: blue, 20 cells from 4 male mice; C57: Cyan, 20 cells from 4 male mice). Data shown as mean ± SEM. Statistical analysis was performed using Two-way RM ANOVA followed by Bonferroni’s multiple comparisons test. **(t)** The maximum action potential number of the MPOA^Esr1^ cells as shown in **(s)**. Data shown as mean ± SEM. Kruskal-Wallis test followed with Two-stage linear step-up procedure of Benjamini, Krieger and Yekutieli. **(u)** *F–I* curves of BNSTp^Esr1^ cells from SW hostile virgin males (red, 20 cells from 4 male mice) and females (gray, 36 cells from 3 female mice). Data shown as mean ± SEM. Statistical analysis was performed using Two-way RM ANOVA followed by Bonferroni’s multiple comparisons test. **(v)** The maximum action potential number of BNSTp^Esr1^ cells as shown in (**u**). Data shown as mean ± SEM. Mann-Whitney test. **(w)** *F–I* curves of BNSTp^Esr1^ cells from SW fathers (blue, 20 cells from 4 male mice) and mothers (gray, 51 cells from 9 female mice). Data shown as mean ± SEM. Statistical analysis was performed using Two-way RM ANOVA followed by Bonferroni’s multiple comparisons test. **(x)** The maximum action potential number of BNSTp^Esr1^ cells as shown in (**w**). Data shown as mean ± SEM. Mann-Whitney test. **(y)** *F–I* curves of MPOA^Esr1^ cells from SW hostile virgin males (red, 20 cells from 4 male mice) and females (gray, 25 cells from 3 female mice). Data shown as mean ± SEM. Statistical analysis was performed using two-way RM ANOVA followed by Bonferroni’s multiple comparisons test. **(z)** The maximum action potential number of MPOA^Esr1^ cells as shown in (**y**). Data shown as mean ± SEM. Mann-Whitney test. **(aa)** *F–I* curves of MPOA^Esr1^ cells from SW fathers (blue, 20 cells from 4 male mice) and mothers (gray, 26 cells from 3 female mice). Data shown as mean ± SEM. Statistical analysis was performed using two-way RM ANOVA followed by Bonferroni’s multiple comparisons test. (**bb**) The maximum number of action potentials of MPOA^Esr1^ cells as shown in (**aa**). Data shown as mean ± SEM. Two-tailed Unpaired t-test.

We observed distinct, parental state-dependent changes in the excitability of MPOA^Esr1^ and BNSTp^Esr1^ cells (**Fig. 6g-l**). In hostile virgin SW males, MPOA^Esr1^ cells displayed a tendency toward depolarization block, with firing rates peaking at approximately 80 pA of current injection and failing to sustain high-frequency spiking with increased current (**Fig. 7h**). In contrast, MPOA^Esr1^ cells in fathers exhibited continuous increases in firing rate with larger current injections—up to ∼180 pA—and maintained high spiking activity at higher current levels (**Fig. 7h**). The maximum firing rate of MPOA^Esr1^ cells in fathers is significantly higher than that in hostile virgins (**Fig. 7i**).

Conversely, BNSTp^Esr1^ cells in hostile virgin males were more excitable than those in fathers. In hostile virgin SW males, BNSTp^Esr1^ cell firing rates continued to increase and remained at a high level with further increased currents. In fathers, however, excitability of BNSTp^Esr1^ cells began to decline after 80 pA of injected current (**Fig. 7j-l**). Overall, the opposite change in excitability—MPOA^Esr1^ cells becoming more excitable and BNSTp^Esr1^ cells becoming less excitable in fathers—likely underlies the altered *in vivo* response patterns to pups observed during the transition to fatherhood.

We further compared the excitability of MPOA^Esr1^ and BNSTp^Esr1^ cells directly (**Fig. 7m-p**). In hostile virgin SW males, BNSTp^Esr1^ cells showed significantly higher excitability than MPOA^Esr1^ cells (**Fig. 7m-n**). In contrast, in SW fathers, MPOA^Esr1^ cells were significantly more excitable than BNSTp^Esr1^ cells (**Fig. 7o-p**). This reversal in excitability balance between MPOA^Esr1^ and BNSTp^Esr1^ cells is likely key to the behavioral switch during fatherhood.

In females, BNSTp^Esr1^ cells in C57 females show significantly lower excitability than SW females^14^. Over 50% of BNSTp^Esr1^ cells in female C57 mice could not fire more than two spikes regardless of the amount of injected currents^14^. Here, we additionally recorded BNSTp^Esr1^ cells in C57 hostile virgin males and fathers and found no significant difference in BNSTp^Esr1^ cell excitability between C57 and SW males (**Fig. 7q-t**). As in the case of SW males, BNSTp^Esr1^ cells in C57 males showed lower excitability in fathers than in hostile virgins (**Fig. 7q-r**). In both strains, all recorded BNSTp^Esr1^ cells were capable of firing more than two spikes upon current injection. MPOA^Esr1^ cell excitability was also similar between C57 and SW males: higher in fathers and lower in naïve hostile males (**Fig. 7s-t**). The similar excitability of BNSTp^Esr1^ cells, as well as MPOA^Esr1^ cells, between C57 and SW males is consistent with the fact that infanticide tendency is high in both C57 and SW males.

Virgin male mice typically show a higher rate of infanticide than virgin females^20^, whereas mothers exhibit more robust parental behaviors than fathers^29–31^. We wondered whether these sex differences in pup-directed behaviors can be explained by the excitability of MPOA^Esr1^ and BNSTp^Esr1^ cells. We compared BNSTp^Esr1^ and MPOA^Esr1^ cell frequency-current (F-I) curves between hostile virgin SW males and females, and between SW fathers and mothers (female data from reference^14^). We found that BNSTp^Esr1^ cells in males exhibited significantly higher firing frequencies than those in females, regardless of reproductive state (**Fig. 6u-x**), whereas MPOA^Esr1^ cells were more excitable and could reach higher firing rates in females than in males, also independent of reproductive state (**Fig. 6y-bb**). Thus, the balance between MPOA^Esr1^ and BNSTp^Esr1^ excitability can be influenced by sex, genetic background, and reproductive state, ultimately determining the pup-directed behaviors of each individual.

## Discussion

Our current and recent studies demonstrated a central role of BNSTp^Esr1^ cells in male and female infanticide^14^. In both sexes, BNSTp^Esr1^ cells are highly activated during infanticide but not parental behaviors, and can bi-directionally control pup-directed attack. In contrast, MPOA^Esr1^ cells in both males and females show high activity during parental behaviors but not infanticide and suppress infanticide. Furthermore, BNSTp^Esr1^ and MPOA^Esr1^ cells form a seesaw relationship through mutual inhibition. During parenthood, BNSTp^Esr1^ cell excitability decreases while MPOA^Esr1^ cell excitability increases to mediate the switch from negative to positive pup-directed behaviors in both mothers and fathers. These results suggest that the same BNSTp^Esr1^-MPOA^Esr1^ circuit controls behaviors towards the young in both sexes. However, the “set point” of MPOA^Esr1^ and BNSTp^Esr1^ cell properties differs between sexes, which likely governs the different tendencies of males and females in expressing pup care versus killing.

### BNSTp^Esr1^ function in infanticide and other male social behaviors

Our study demonstrated that BNSTp^Esr1^ cells are indispensable for infanticide, adding to an increasing list of behavioral functions involving BNSTp in males. Previous studies showed that BNSTp^Esr1^ or its subpopulations are essential for male territorial aggression^18,24,32^ and sexual behavior^32–34^. Inhibiting BNSTp Esr1 cells^24^, or their subsets that express aromatase^32^ or Tac1^18,34^, all caused reduced inter-male aggression and male sexual behaviors. In a study focusing specifically on male sexual behaviors, Esr2-expressing cells (also a subset of Esr1 cells) were found to be uniquely and dramatically excited during ejaculation, causing post-ejaculation sexual satiety^33^.

What is the relationship of BNSTp cells involved in different male social functions? Single-cell miniscope calcium imaging revealed that largely distinct BNSTp^Esr1^ cells are excited during adult male-male and male-female interactions^24^. Single-nucleus RNA sequencing (snRNAseq) identified dozens of distinct clusters in the BNSTp^18,35,36^. Thus, male mating and fighting likely involve largely non-overlapping and molecularly differentiable BNSTp subpopulations. Whether infanticide recruits a subpopulation of BNSTp^Esr1^ cells distinct from those activated during male mating and fighting remains to be investigated.

Although multiple loss-of-function studies (including ours) showed an essential role of BNSTp^Esr1^ cells or their subsets in inter-male aggression, activating Esr1, Tac1, or aromatase cells in the BNSTp all failed to induce attack towards adult males^24,32,34^. It is not yet clear whether the failed light-induced attack is due to the co-activation of other social behavior-relevant cells that trumps the effect of aggression cell activation, or BNSTp modulates but is incapable of directly driving attack towards adult males.

In our experiments, BNSTp^Esr1^ cell stimulation consistently evokes allogrooming of adult intruders in both males and females^14^. Recent studies have shown that allogrooming can carry multiple meanings, depending on the context. For instance, MeA^Tac1∩Vgat^ cells were suggested to drive allogrooming to comfort stressed cage mates^37^. In contrast, when directed toward unconscious conspecifics, allogrooming—especially when accompanied by more intense facial nibbling, a behavior we also observed during BNSTp^Esr1^ activation—may represent a “first-aid-like” behavior, aiming at reviving the unresponsive individual^38–40^. The precise meaning of allogrooming induced by BNSTp^Esr1^ cell activation remains unclear, but it may suggest another previously unrecognized social behavior involving BNSTp^Esr1^ cells.

Clearly, male BNSTp^Esr1^ cells are engaged in multiple social behaviors. Nevertheless, when BNSTp^Esr1^ cells are activated as a whole, infanticide is induced robustly in the presence of pups. As we previously proposed^30^, social behavior circuits are organized in layers, with limbic regions serving as “sensory and internal state detectors” to provide permission to the midbrain premotor cells to act upon “immediate releasing cues.” We speculate that when all BNSTp^Esr1^ cells are artificially activated, permissions for multiple social behaviors are granted, and the final motor output is selected based on the immediate releasing cues. When stimulation occurs in the presence of pups, infanticide, not other adult-directed behaviors, is expressed.

### The antagonistic relationship between infanticide and parental behavior circuits

It is well established that MPOA^Esr1^ cells are critical for parental behaviors in females^7–9,27^. A previous study showed that activating MPOA^Esr1^ cells can also induce time-locked pup retrieval in males^27^. Here, we demonstrated that male MPOA^Esr1^ cells are naturally activated during pup caring but not infanticide, further supporting the important role of MPOA^Esr1^ cells in parental behaviors in males. It is worth noting that MPOA^Esr1^ cells, critical for parental care, are likely GABAergic, as broadly activating MPOA GABAergic cells also promotes pup retrieval, whereas activating glutamatergic cells suppresses pup retrieval and induces anxiety^41^.

In both males and females, MPOA^Esr1^ and BNSTp^Esr1^ cells antagonize each other through reciprocal inhibitory synaptic connections^14^. Therefore, reducing the activity of one population not only impairs its mediated behavior but also facilitates behavior driven by the other population. Indeed, in both sexes, inhibiting MPOA^Esr1^ cells not only impairs parental behavior but also facilitates infanticide, whereas impairing BNSTp^Esr1^ cells not only impairs infanticide but also promotes parental behaviors^14^. Such a seesaw organization ensures that positive or negative pup-directed behaviors, but not both, are expressed at a given time.

It is worth noting that BNSTp and MPOA cells do not always antagonize each other at the functional level. In the context of male sexual behavior, both BNSTp Tac1-and MPOA Tacr1 positive cells, which both co-express Esr1, promote mating^34^. In this case, BNSTp^Tac1^ cells are upstream of MPOA^Tacr1^ cells, as the behavior changes induced by BNSTp^Tac1^ manipulation can be overridden by simultaneous opposing manipulation of MPOAp^Tacr1^ cells. The positive “message” sent from BNSTp^Tac1^ to MPOAp^Tacr1^ is through Tacr1 activation, which results in potentiated MPOA responses to excitatory inputs, likely originating from a region outside of BNSTp^34^. Given that BNSTp contains abundant and diverse neuropeptides, the exact influence of BNSTp on downstream cells is likely a combined effect of fast neurotransmitter (mainly GABA) and slower neuropeptide-mediated modulation.

### Neural plasticity of paternal and infanticide circuits during parenthood

Parenthood is associated with dramatic behavioral changes in both females and males^2–6,42–45^. Sexually naïve mice typically ignore or even attack young, but upon becoming parents, they exhibit pronounced shifts toward parenting behaviors. These behavioral transformations must be supported by changes in the underlying neural circuits. A previous study revealed that oxytocin neurons receive increased inputs from the lateral hypothalamus in fathers^46^. Here, we found that the excitability of Esr1-expressing neurons in the BNSTp or MPOA shifts in opposite directions during fatherhood: MPOA^Esr1^ neurons increase their excitability, while BNSTp^Esr1^ neurons show a decrease^14^. These synaptic and cellular changes collectively cause the switch from hostile to caring behaviors during parenthood.

In females, numerous studies have shown that pregnancy triggers profound hormonal changes, driving neural remodeling such as changes in cell morphology, gene expression, and neural excitability^42,47–51^. For example, Rachida et al. showed that estradiol and progesterone remodel MPOA^Gal^ neurons by increasing their excitability, promoting dendritic spine formation, and enhancing excitatory synaptic input during pregnancy^51^. Intriguingly, similar to MPOA cells, BNSTp^Esr1^ cells also express Esr1 and the progesterone receptor (PR) but exhibit excitability changes opposite to those of MPOA^Esr1^ cells. We speculate that the distinct cellular changes in BNSTp and MPOA cells may involve other sex hormone receptors that are differentially expressed in these two regions. For example, Esr2, which has been proposed to oppose Esr1 functions, is more abundantly expressed in the BNSTp than MPOA^18,52^.

In contrast to females, males do not experience a significant hormonal surge during the transition to fatherhood. Only a slight decrease in testosterone and estradiol was reported^53,54^. Yet, male BNSTp^Esr1^ and MPOA^Esr1^ cells show similar excitability changes during parenthood as females. It remains unclear whether sex hormones are responsible for the behavior switch in fathers. Intriguingly, paternal behaviors could emerge in male mice 3 weeks after ejaculation, even when they are housed alone, suggesting a slow mechanism triggered by ejaculation is likely responsible for the switch^5,31^. Recently, several studies showed post-ejaculation neural changes in the MPOA and BSNTp that unfold over several days^33,55^. The molecular and cellular processes supporting ejaculation-induced neural and behavioral changes are an active field of investigation.

### Sex differences in the infanticide circuit

Compared to females, naïve laboratory male mice show a higher tendency to attack and a lower tendency to care for pups^20^. This sex difference in pup-directed behaviors is presumably due to differences in the underlying circuits. Indeed, we found that BNSTp^Esr1^ cells are more excitable in males than females, while MPOA^Esr1^ cells are more excitable in females than males, regardless of the animal’s genetic background and reproductive state. Consistent with our findings, a recent study revealed that male MPOA GABAergic cells require higher excitatory inputs to spike^56^. The differences in cellular properties are likely a product of molecular differences. In support, snRNAseq demonstrated an extensive array of differentially expressed genes in the MPOA and BNSTp Esr1-expressing cells between sexes^18^. Anatomically, male BNSTp and MPOA are almost twice the size of females and appear to support more social functions^57^. For example, MPOA is indispensable for male but not female sexual behaviors^27,58,59^; inhibiting BNSTp cells impairs aggressive and sexual behaviors in males but not females^14,24,32^. Thus, it is likely that a smaller fraction of BNSTp^Esr1^ or MPOA^Esr1^ cells is related to pup-directed behaviors in males than in females.

Both male and female BNSTp^Esr1^ and MPOA^Esr1^ cells form mutual inhibition. The inhibition appears to be stronger from BNSTp^Esr1^ to MPOA^Esr1^ cells than from MPOA^Esr1^ to BNSTp^Esr1^ cells in both sexes. Intriguingly, we noted that a higher percentage of BNSTp^Esr1^ cells receive excitatory inputs from MPOA^Esr1^ in males (87%) than in females (29%). The stronger excitatory inputs from MPOA^Esr1^ to BNSTp^Esr1^ may diminish the MPOA inhibitory control on BNSTp, favoring activation of the infanticide circuit.

Our current findings in males, together with our recent study in females^14^, suggest BNSTp^Esr1^-MPOA^Esr1^ circuit is similarly organized to mediate infanticide and parental behaviors in both sexes. Quantitative differences in pup-directed behavior can be attributed to variations in the number of cells, their intrinsic properties, and possibly synaptic strength. We speculate that other regions, e.g., MEApd^12,16^, PFA^12^, and BNSTrh^13^, that were previously found critical for male infanticide likely also play a role in female infanticide, although this needs to be proven experimentally. Regarding other social behaviors, adult-directed attack has also been found to engage a similar set of brain regions (except BNSTp^32^), such as VMHvl^60,61^, PMv^62–64^, MeApd^65,66^, and SI^67^, in both sexes. In contrast, key regions for male and female sexual behaviors are qualitatively different, with MPOA being the most crucial region for males and VMHvl for females^30^. Notably, males and females express aggression or infanticide using a series of qualitatively similar movements, while male and female sexual behaviors differ dramatically in their motor patterns. These results collectively suggest that sex-specific circuits are likely reserved for sex-specific behaviors. Quantitative differences in social behaviors between sexes could be achieved through fine-tuning the sex-common behavior circuits.

## Acknowledgments

We thank all of the members of the Lin laboratory for feedback, and Yiwen Jiang and Prakhar Dua for assisting with genotyping and maintaining the mouse lines. This work was supported by the Fundamental Research Funds for the Central Universities (to L.M. and B.Y.), National Natural Science Foundation of China grants 82501858 and 32571196 (to L.M.), Overseas Excellent Young Scientists Fund (to L.M.), and Levy Leon Postdoctoral Fellowship (to L.M.); the NIH grants R01MH101377, R01MH124927, 1R01HD092596, and U19NS107616 (to D.L.).

## Author contributions

D.L. and L.M. conceived the project, designed experiments, analyzed data, and wrote the paper. D.L. supervised the project. L.M., Y.W., Q.W., and Y.X. performed all the experiments. B.Y. provided critical feedback on the experiments and manuscript.

## MATERIALS AND METHODS

### Mice

All procedures were approved by the NYULMC and WHU Institutional Animal Care and Use Committee (IACUC) in compliance with the National Institutes of Health (NIH) Guidelines for the Care and Use of Laboratory Animals. Adult mice (8-32 weeks) were used as test subjects for all studies. Mice were housed under a reversed 12-hour light-dark cycle (dark cycle, 10 a.m. to 10 p.m.), with food and water available ad libitum. Room temperature was kept between 22 – 26 °C and humidity between 30-70%, with a daily average of approximately 45%. Esr1-2A-Cre mice of C57BL/6 background were purchased from Jackson Laboratory (stock no. 017911). Esr1-2A-Cre mice of SW background were backcrossed with SW wild-type mice for at least four generations. Vgat-Flp mice were purchased from Jackson Laboratory (stock no. 029591). Esr1-2A-Cre::Vgat-Flp mice were generated by crossing SW Esr1-2A-Cre mice with Vgat-Flp mice. Wild-type SW mice were purchased from Taconic. Wildtype Balb/c mice were purchased from Charles River. P1-P5 pups used for behavioral experiments were bred in-house. All mice were group-housed until adulthood. After surgery, mice were singly housed. All experiments were performed during the dark phase of the daily cycle.

### Viruses

AAV2-hSyn-DIO-GFP, AAV5-hSyn-flex-ChrimsonR-tdTomato, and AAV2-EF1a-DIO-ChR2-EYFP were purchased from the University of North Carolina vector core. AAV2-CAG-flex-GCaMP6f was purchased from the University of Pennsylvania vector core. AAV2-hSyn-DIO-hM3Dq-mCherry, AAV2-hSyn-DIO-hM4Di-mCherry, AAV8-nEF-Con/Fon-ChRmine-oScarlet, and AAV2-hSyn-DIO-mCherry were purchased from Addgene. AAV8-hSyn-DIO-DTR was purchased from Boston Children’s Hospital. AAV9-hSyn-DIO-hChR2(H134R)-mCherry-ER2-WPRE-pA were purchased from Taitool. All viruses were stored at -80 °C until use. The titer of each virus ranged from 2 × 10^12^ to 2 × 10^13^ genomic copies per ml.

### Stereotactic Surgery

Mice (8-32 weeks old) were anesthetized with 1%-2% isoflurane and positioned on a stereotaxic apparatus (Kopf Instruments Model 1900). Viruses were injected into the brain using a glass capillary attached to a nanoinjector (World Precision Instruments, Nanoliter 2000).

For fiber photometry recording of BNSTp^Esr1^ activity, 200 nL AAV2-CAG-Flex-GCaMP6f was unilaterally injected into the BNSTp (AP: -0.45 mm, ML: -0.9 mm, DV: -3.6 mm) of heterozygous virgin Esr1-2A-Cre SW male mice. For fiber photometry recording of MPOA^Esr1^ activity, 200nL AAV2-CAG-Flex-GCaMP6f was unilaterally injected into the MPOA (AP: 0 mm, ML: -0.3 mm, DV: -4.95 mm) of heterozygous virgin Esr1-2A-Cre males. A 400-µm optical fiber assembly (Thorlabs, FR400URT, CF440) was inserted 400 µm above the virus injection site and secured on the skull using adhesive dental cement (C&B Metabond, S380) after the virus injection. All recordings started approximately 3 weeks after surgery. All male mice were screened before surgery. During the screening, 2-3 pups were introduced into the testing mouse home cage and tested for around 5 min. Only male mice that showed spontaneous infanticide were used.

To optogenetically activate BNSTp^Esr1^ neurons, 200nL AAV2-EF1a-DIO-ChR2-EYFP (control: AAV2-hSyn-DIO-GFP) was bilaterally injected into the BNSTp (AP: - 0.45 mm, ML: ±0.9 mm, DV: -3.6 mm) of adult Esr1-2A-Cre SW male mice. Two 200-µm optical fiber assemblies (Thorlabs, FT200EMT, CFLC230) were implanted 400 µm above the virus injection sites and secured on the skull using adhesive dental cement (C&B Metabond, S380) after virus injection. All male mice were screened prior to surgery, and only male mice that did not show spontaneous infanticide were used.

To chemogenetically activate BNSTp^Esr1^ neurons, 200nL AAV2-hSyn-DIO-hM3Dq-mCherry was bilaterally injected into the BNSTp (AP: -0.45 mm, ML: ±0.9 mm, DV: -3.6 mm) of adult Esr1-2A-Cre SW males. All male mice were screened before surgery, and only males that did not show spontaneous infanticide were used.

To chemogenetically inhibit BNSTp^Esr1^ neurons, 200nL AAV2-hSyn-DIO-hM4Di-mCherry (control: AAV2-hSyn-DIO-mCherry) was bilaterally injected into the BNSTp (AP: -0.45 mm, ML: ±0.9 mm, DV: -3.6 mm) of adult Esr1-2A-Cre C57BL/6 males. All male mice were screened before surgery, and only males that showed spontaneous infanticide were used.

To ablate BNSTp^Esr1^ cells, 200nL AAV8-hSyn-DIO-DTR (control: AAV2-hSyn-DIO-mCherry) was bilaterally injected into the BNSTp (AP: -0.45 mm, ML: ±0.9 mm, DV: -3.6 mm) of adult Esr1-2A-Cre SW males. All male mice were screened before surgery, and only males that showed spontaneous infanticide were used.

To optogenetically activate BNSTp^Esr1^ projection to the MPOA and simultaneously record MPOA^Esr1^ cell activity, 200 nL AAV5-hSyn-flex-ChrimsonR-tdTomato was injected into the BNSTp unilaterally (AP: -0.45 mm, ML: ±0.9 mm, DV: -3.6 mm) and at the same time 200 nL AAV2-CAG-Flex-GCaMP6f was injected into the ipsilateral MPOA (AP: 0 mm, ML: -0.3 mm, DV: -4.95 mm) of adult Esr1-2A-Cre SW male and female mice. After virus injection, a 400-µm optical fiber assembly (Thorlabs, FR400URT, CF440) was implanted 400 µm above the MPOA (AP: 0 mm, ML: -0.3 mm, DV: -4.55 mm) and secured on the skull using adhesive dental cement (C&B Metabond, S380). The recording started at least 4 weeks after surgery.

To optogenetically activate MPOA^Esr1∩Vgat^ projection to the BNSTp and simultaneously record MPOA^Esr1^ cell activity, 200 nL AAV8-nEF-Con/Fon-ChRmine-oScarlet was injected into the MPOA unilaterally (AP: 0 mm, ML: -0.3 mm, DV: -4.95 mm) and at the same time 200 nL AAV2-CAG-Flex-GCaMP6f was injected into the ipsilateral BNSTp (AP: -0.45 mm, ML: ±0.9 mm, DV: -3.6 mm) of adult Esr1-2A-Cre::Vgat-Flp male and female mice. After virus injection, a 400-µm optical fiber assembly (Thorlabs, FR400URT, CF440) was implanted 400 µm above the BNSTp (AP: -0.45 mm, ML: ±0.9 mm, DV: -3.6 mm) and secured on the skull using adhesive dental cement (C&B Metabond, S380). The recording started at least 4 weeks after surgery.

To chemogenetically inhibit MPOA^Esr1^ neurons, 300 nL AAV2-hSyn-DIO-hM4Di-mCherry was bilaterally injected into the MPOA (AP: 0 mm, ML: -0.3 mm, DV: -4.95 mm) of Esr1-2A-Cre male mice. All male mice were screened before surgery, and only males that did not show spontaneous infanticide were used.

To chemogenetically activate MPOA^Esr1^ neurons, 300 nL AAV2-hSyn-DIO-hM3Dq-mCherry was bilaterally injected into the MPOA (AP: 0 mm, ML: -0.3 mm, DV: -4.95 mm) of Esr1-2A-Cre male mice. All male mice were screened before surgery, and only males that showed spontaneous infanticide were used.

For examining the synaptic connection from BNSTp^Esr1^ cells to MPOA^Esr1^ cells, AAV9-hSyn-DIO-hChR2(H134R)-mCherry-ER2-WPRE-pA was bilaterally injected into BNSTp (AP: -0.45 mm, ML: ±0.9 mm, DV: -3.6 mm; 300 nl/side), and at the same time AAV2-hSyn-DIO-GFP was injected bilaterally into MPOA (AP: 0 mm, ML: ±0.3 mm, DV: -4.95 mm; 300 nl/side). For MPOA^Esr1^ to BNSTp^Esr1^ projection, AAV9- hSyn-DIO-hChR2(H134R)-mCherry-ER2-WPRE-pA was bilaterally injected into MPOA (AP: 0 mm, ML: ±0.3 mm, DV: -4.95 mm; 300 nl/side), and at the same time AAV2-hSyn-DIO-GFP was bilaterally injected into BNSTp (AP: -0.45 mm, ML: ±0.9 mm, DV: -3.6 mm; 300 nl/side). All mice were heterozygous virgin Esr1-2A-Cre C57BL/6 male mice.

For examining the intrinsic properties of MPOA^Esr1^ and BNSTp^Esr1^ cells, AAV2-hSyn-FLEX-GFP virus was bilaterally injected into MPOA (AP: 0 mm, ML: ±0.3 mm, DV: -4.95 mm; 300 nl/side) and BNSTp (AP: -0.45 mm, ML: ±0.9 mm, DV: -3.6 mm; 300 nl/side) in the same animal. All mice were heterozygous virgin Esr1-2A-Cre male mice in SW or C57BL/6 background, and were screened both prior to surgery and again on the day before brain slice collection for electrophysiological recording.

### Fiber photometry

For fiber photometry recording, the setup was the same as described previously^14^. Briefly, a bandpass filtered (passing band: 472 ± 15 nm, FF02-472/30-25, Semrock) 390-Hz sinusoidal blue LED light (30 µW; LED light: M470F1; LED driver: LEDD1B; from Thorlabs) was used to excite GCaMP6f through a 400 µm optic fiber. The emission light passed through the same optic fiber was bandpass filtered (passing bands: 535 ± 25 nm, FF01-535/505, Semrock), went through an adjustable zooming lens (Thorlab, SM1NR01 and Edmund optics, #62-561), was detected by a Femtowatt Silicon Photoreceiver (Newport, 2151) and recorded with a real-time processor (RP2, TDT).

For fiber photometry recording of BNSTp^Esr1^ and MPOA^Esr1^ neurons, sexually naïve Esr1-2A-Cre male mice were injected with AAV2-CAG-Flex-GCaMP6f into the BNSTp and MPOA, respectively. Mice were single-housed after surgery. The recording started 3 weeks after surgery. During the first recording section, a P1-P5 pup was introduced into the recording animal’s home cage at a location away from the nest. If and once infanticide occurred, we removed and euthanized the pup. A total of 3-5 pups were introduced during the recording session, each for approximately 1-2 minutes. For BNSTp^Esr1^ cell recording, after the first recording session with pups, an adult non-aggressive BALB/c male mouse was introduced into the recording animal’s home cage, and then recorded 10 minutes. After recording in naïve animals, each male was paired with a female mouse. 3-4 days after the female gave birth, all male mice were recorded again. During the recording session, the female and pups were removed from the cage 10 minutes before the recording started, and then a total of 6-9 of the male’s own pups were introduced into the recording animal’s home cage sequentially at a location distant from the nest. Retrieved pups were kept in the nest without being disturbed during the recording.

For fiber photometry recording of MPOA^Esr1^ neurons while simultaneously optogenetically activating BNSTp^Esr1^ cells, Esr1-2A-Cre males and females were injected with AAV2-CAG-Flex-GCaMP6f into the MPOA and AAV5-hSyn-flex-ChrimsonR-tdTomato into the BNSTp. For fiber photometry recording of BNSTp^Esr1^ neurons while simultaneously optogenetically activating MPOA^Esr1∩Vgat^ cells, Esr1-2A-Cre::Vgat-Flp males and females were injected with AAV2-CAG-Flex-GCaMP6f into the BNSTp and AAV8-nEF-Con/Fon-ChRmine-oScarlet into the MPOA. All recordings started four weeks post-surgery. During the recording, animals were freely moving in their home cages. Stimulation consisted of five sham trials (0 mW light) and five 589 nm yellow light trials (3 mW, 20 Hz, 20 ms pulses, 20 s per trial).

To analyze the recording data, the MATLAB function ‘‘msbackadj’’ with a moving window of 1/8 of the total recording duration was first applied to obtain the moment-to-moment baseline signal (F_baseline_). The ΔF/F was then calculated as (F_raw_ – F_baseline_)/F_baseline_. The Z-Scored ΔF/F then was calculated as (ΔF/F – mean (ΔF/F))/STD(ΔF/F). AUC was calculated as the area under the curve using MATLAB function “trapz” divided by the accumulated time of the behavior. The PETHs were constructed by aligning the Z-Scored ΔF/F to the onset of each trial of a behavior, averaging across all trials for each animal, and then averaging across animals.

### Optogenetic activation of BNSTp^Esr1^ cells

For optogenetic activation of BNSTp^Esr1^ neurons, non-infanticidal Esr1-2A-Cre male mice were injected with AAV2-EF1a-DIO-ChR2-EYFP (Control: AAV2-hSyn-DIO-GFP) into the BNSTp. Three weeks after surgery, we delivered 20Hz 473 nm blue laser pulses through the implanted optic fiber. Each test session began with 2-4 times of sham (laser off) stimulations, followed by interleaved light stimulation (0.5-2 mW) and sham stimulation. Each stimulation started when male mice investigated a pup or an adult BALB/c male. Once the male mice attacked and wounded the pup, the pup was quickly removed, euthanized, and replaced with a new pup.

### BNSTp^Esr1^ cell ablation

SW males were prescreened, and only males that showed stable infanticide behavior were used for surgery. During the screening, 1-2 pups were introduced into the home cage of the test male mouse for 10 minutes on three consecutive days. 4 weeks after surgery, we tested the test mouse’s behaviors towards the pups the day before DT injection. During the test, we introduced 1-2 P1-P5 pups into the test male’s home cage at a location away from the nest for 10 minutes or until infanticide occurred, whichever came first. After the test, we injected DT (50 µg/kg, 5 μg/ml dissolved in PBS) intraperitoneally into each male. 8 days later, males were tested again on three consecutive days by introducing 1-2 P1-P5 pups into the test male’s home cage at a location away from the nest for 10 minutes. If male mice attacked and wounded pups, the pups were euthanized immediately after the test.

### Chemogenetic inhibition and activation

For chemogenetic inhibition of BNSTp^Esr1^ neurons, AAV2-hSyn-DIO-hM4Di-mCherry (Control: AAV2-hSyn-DIO-mCherry) was bilaterally injected into BNSTp of C57 Esr1-2A-Cre male mice. For chemogenetic activation of BNSTp^Esr1^ neurons, AAV2-hSyn-DIO-hM3Dq-mCherry was bilaterally injected into BNSTp of SW Esr1-2A-Cre male mice. For chemogenetic inhibition and activation of MPOA^Esr1^ neurons, AAV2-hSyn-DIO-hM4Di-mCherry and AAV2-hSyn-DIO-hM3Dq-mCherry were bilaterally injected into the MPOA of SW Esr1-2A-Cre non-infanticidal male mice, respectively.

All animals were first tested after saline injection and then after CNO (1 mg/kg) injection. For the pup-directed behavior test, 30 minutes after saline or CNO injection, we scattered 1-3 P1-P5 pups in the test male’s home cage for 10 minutes or until the infanticide occurred, whichever came first. If the male mouse attacked and injured the pup, the wounded pup was quickly removed, euthanized, and the recording was terminated. For BNSTp^Esr1^ inactivation and activation experiments, we also tested the behaviors against an adult male intruder. 30 minutes after saline or CNO injection, we introduced a non-aggressive adult BALB/c male into the test male’s home cage for 10 minutes.

### In vitro electrophysiological recording

For *in vitro* whole-cell patch-clamp recordings, mice were anaesthetized with isoflurane, and the brains were removed and submerged in oxygenated ice-cold cutting solution containing 110 mM choline chloride, 25 mM NaHCO_3_, 2.5 mM KCl, 7 mM MgCl_2_, 0.5 mM CaCl_2_, 1.25 mM NaH_2_PO_4_, 25 mM glucose, 11.6 mM ascorbic acid, and 3.1 mM pyruvic acid. The coronal MPOA and BNSTp 275 µm brain sections were cut using the Leica VT1200s vibratome and incubated in artificial cerebral spinal fluid containing 125 mM NaCl, 2.5 mM KCl, 1.25 mM NaH_2_PO_4_, 25 mM NaHCO_3_, 1 mM MgCl_2_, 2 mM CaCl_2_ and 11 mM glucose at 34 °C for 30 min and then at room temperature until use. Next, we moved MPOA or BNST containing sections into the recording chamber perfused with oxygenated ACSF and performed the whole-cell recordings. The recorded signals were acquired with MultiClamp 700B amplifier (Molecular Devices) and digitized at 20 kHz using DigiData1550B (Molecular Devices). The stimulation and recording were conducted using the Clampex 11.0 software (Axon Instruments). The intracellular solution for current-clamp recording contained 145 mM K-gluconate, 2 mM MgCl_2_, 2 mM Na_2_ATP, 10 mM HEPES, 0.2 mM EGTA (286 mOsm, pH 7.2). The recorded electrophysiological data were analyzed using Clampfit (Molecular Devices).

The synaptic transmission between BNSTp^Esr1^ and MPOA^Esr1^ cells was assessed by voltage-clamping postsynaptic cells at 0 mV and -70 mV to record optogenetically evoked IPSCs and EPSCs, respectively. Pipettes were filled with internal solution containing (in mM): 135 CsMeSO₃, 10 HEPES, 1 EGTA, 3.3 QX-314 chloride, 4 Mg-ATP, 0.3 Na-GTP, and 8 sodium phosphocreatine (290–300 mOsm, pH 7.2 with CsOH). ChR2-expressing axons were stimulated with ten blue light pulses (0.5 ms, 0.1 Hz). TTX (1 μM), 4-AP (100 μM), and picrotoxin (100 μM) were bath-applied sequentially for 10–20 min each.

To determine the intrinsic excitability of these cells, we labeled Esr1 cells in the BNSTp and MPOA with mCherry and identified them with an Olympus 40× water-immersion objective with an MDF-MCHC filter (THORLABS). We performed current-clamp recordings and injected current steps ranging from −20 pA to 260 pA in 20 pA increments into the recorded cell. The total number of spikes during each 500-ms-long current step was then used to construct the *F*–*I* curve.

### Behavioral analyses

Behaviors were analyzed frame-by-frame using custom software written in MATLAB (https://pdollar.github.io/toolbox/). Investigating pup is defined as when the animal’s nose closely contacts any body parts of a pup. Attacking pup is defined as biting a pup and is confirmed by the wounds. Retrieving pup is from the moment when the animal lifts the pup using its jaw to the moment when the pup is dropped in or around the nest. Investigating male is defined as nose-to-face, nose-to-trunk, or nose-to-urogenital contact with an intruder male. Attacking male is defined as lunging, biting, and fast movements connecting these behaviors. Grooming male is defined as licking or grooming the head or neck area of the adult male intruder.

### Histological analysis

Mice were first perfused with 1 × PBS, followed by 4% PFA. Brains were dissected, post-fixed in 4% PFA overnight at 4℃, rinsed with 1 × PBS, and dehydrated in 30% sucrose for 12-16 hours. 30 µm sections were cut on a Leica CM1950 cryostat. For Esr1 staining of BNSTp^Esr1^-DTR animals, every third brain section was collected. Then, free-floating brain slices were rinsed with PBS (3 × 10 min) and PBST (0.1% Triton X-100 in PBS, 1 × 30 minutes) at room temperature, followed by 1 hour of blocking in 10% normal donkey serum at room temperature. The primary antibody (Rabbit anti-Esr1, 1:1000 dilution, Invitrogen, Cat. # PA1-309) was diluted in PBST with 3% normal donkey serum and incubated overnight (12-16 hours) at 4 °C. Brain slices were then washed with PBST (3 × 10 min) and incubated with the secondary antibody (Secondary antibody for Esr1 staining: 488-Donkey anti-rabbit, 1:500 dilution, Jackson ImmunoResearch, Cat. # 711-685-152) for 2 hours at room temperature. Then, brain slices were washed with PBST (3 × 10 min), rinsed with 1 × PBS, and mounted on slides (Fisher Scientific, 12-550-15), dried 10 min at room temperature, and coverslipped using 50% glycerol containing DAPI (Invitrogen, Cat. #00-4959-52). Images were acquired using a slide scanner (Olympus, VS120) or a confocal microscope (Zeiss LSM 700 microscope). Brain regions were identified based on the Allen Mouse Brain Atlas, and cells were counted manually using ImageJ.

### Statistics

All statistical analyses were performed using MATLAB R2023a and Prism 10 software. McNemar’s test was used to analyze paired nominal data from two groups. To compare the mean of two groups, paired t-test (for paired data) or unpaired t-test (for unpaired data) was used if both samples passed Kolmogorov – Smirnov tests for normality. Otherwise, Wilcoxon matched-pairs signed rank test (for paired data) or Mann Whitney test (for unpaired data) was used. For comparisons of mean values of more than two groups, RM one-way ANOVA followed by Bonferroni’s multiple comparison tests was used if all samples passed Kolmogorov – Smirnov tests for normality. Otherwise, Kruskal-Wallis test followed with Two-stage linear step-up procedure of Benjamini, Krieger and Yekutieli. For comparisons of more than two variables across multiple groups, we used two-way ANOVA. Specifically, the mixed-effects model was used when comparing between-subject and within-subject variables. Two-way RM ANOVA was used when performing within-subject comparisons for both variables. Two-way AVNOV tests were followed by Bonferroni’s multiple comparison tests. All statistical tests are two-tailed. For all statistical tests, p values below 0.1 are indicated in the figures. All error bars are shown as SEM.

---

**Figure S1.**
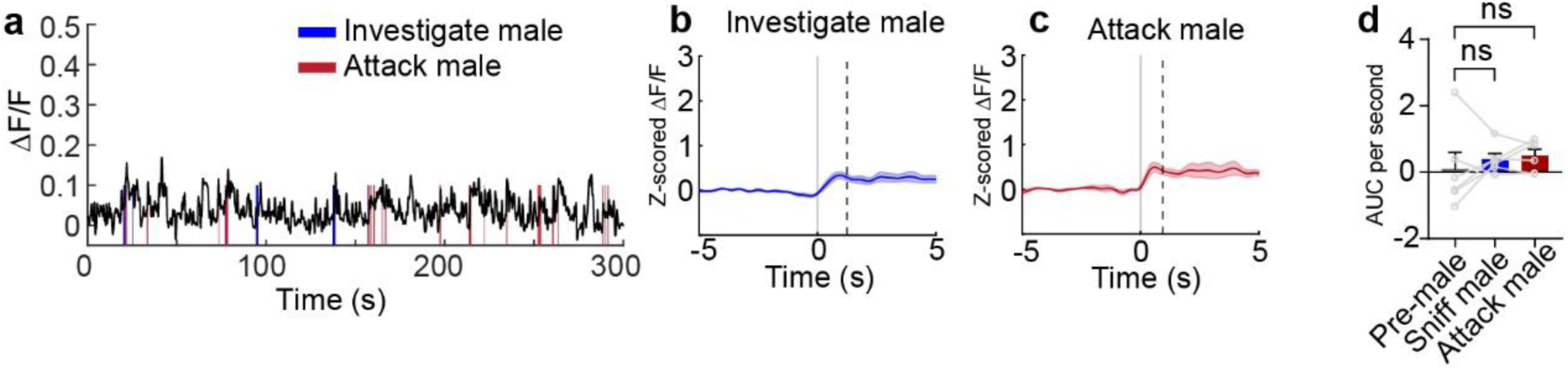
The response pattern of BNSTp^Esr1^ cells during territory aggression in SW male mice. **(a)** Representative ΔF/F traces of BNSTp^Esr1^ cells during territory aggression. **(b and c)** PETHs of Z-scored ΔF/F of BNSTp^Esr1^ cells aligned to the onset of investigating (**b**) and attacking a male intruder (**c**). (n=6 mice, data shown as mean ± SEM) **(c)** The average area under the curve (AUC) of Z-scored ΔF/F per second during various intruder male-directed behaviors to compare responses across behaviors. (n=6 male mice. RM one-way ANOVA followed by Bonferroni’s multiple comparisons test; ns: not significant. Data shown as mean + SEM)

**Figure. S2.**
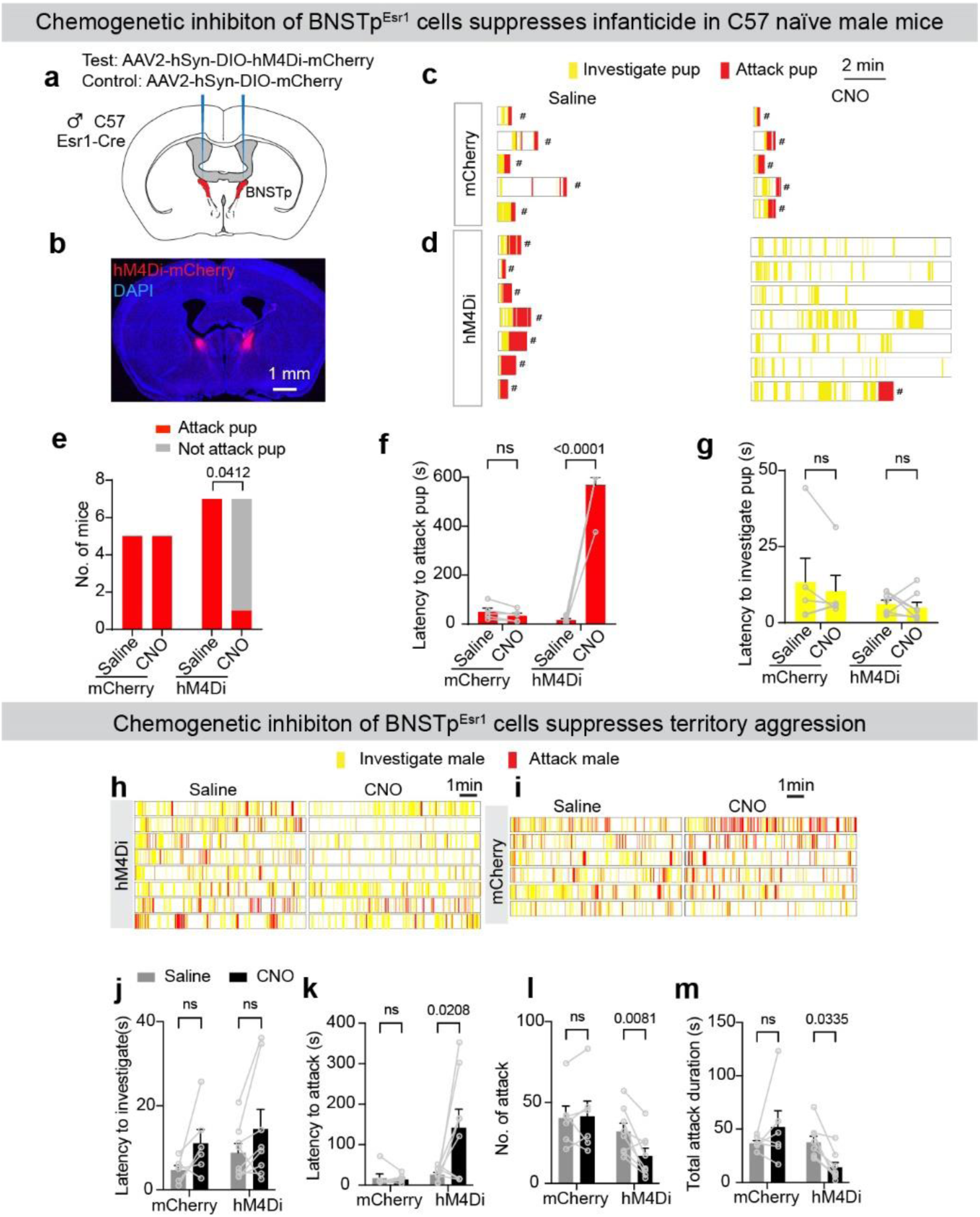
Chemogenetic inhibition of BNSTp^Esr1^ neurons suppresses infanticide and territorial aggression. **(a)** The strategy to chemogenetically inhibit BNSTp^Esr1^ cells. **(b)** A representative image showing hM4Di-mCherry (red) expression in BNSTp. **(c and d)** Raster plots showing pup-directed behaviors in mCherry (**c**) and hM4Di (**d**) male mice after saline (left) and CNO (right) injection. # wounded pups were removed and euthanized. **(d)** The number of mCherry and hM4Di male mice that attacked and did not attack pups after saline and CNO injection. (McNemar’s test; ns: not significant; p=0.0412) **(e)** Latency to attack pups in mCherry and hM4Di male mice after saline and CNO injection. The latency equals 600 s if no pup attack occurs during the 10 minutes of the test. (n = 5 males of mCherry group, n = 7 males of hM4Di group. Mixed effect two-way ANOVA followed by Bonferroni’s multiple comparisons test; ns: not significant. Data shown as mean + SEM.) **(f)** Latency to investigate pups in mCherry and hM4Di male mice after saline and CNO injection. (n = 5 males of mCherry group, n = 7 males of hM4Di group. Mixed effect two-way ANOVA followed by Bonferroni’s multiple comparisons test, ns: not significant. Data shown as mean + SEM) **(h and i)** Raster plots showing adult male intruder-directed behaviors in hM4Di (**h**) and mCherry (**i**) male mice after saline (left) and CNO (right) injection. **(j)** Latency to investigate adult male intruders in mCherry and hM4Di male mice after saline and CNO injection. (n = 6 males of mCherry group, n = 8 males of hM4Di group. Mixed effect two-way ANOVA followed by Bonferroni’s multiple comparisons test; ns: not significant; Data shown as mean + SEM.) **(k)** Latency to attack adult male intruders in mCherry and hM4Di male mice after saline and CNO injection. (n = 6 males of mCherry group, n = 8 males of hM4Di group. Mixed effect two-way ANOVA followed by Bonferroni’s multiple comparisons test; ns: not significant. Data shown as mean + SEM.) **(l)** Number of attacks towards adult male intruders in mCherry and hM4Di male mice after saline and CNO injection. (n = 6 males of mCherry group, n = 8 males of hM4Di group. Mixed effect two-way ANOVA followed by Bonferroni’s multiple comparisons test; ns: not significant. Data shown as mean + SEM) **(m)** Total attack duration towards adult male intruders in mCherry and hM4Di male mice after saline and CNO injection. (n = 6 males of mCherry group, n = 8 males of hM4Di group. Mixed effect two-way ANOVA followed by Bonferroni’s multiple comparisons test; ns: not significant. Data shown as mean + SEM.)

**Figure. S3.**
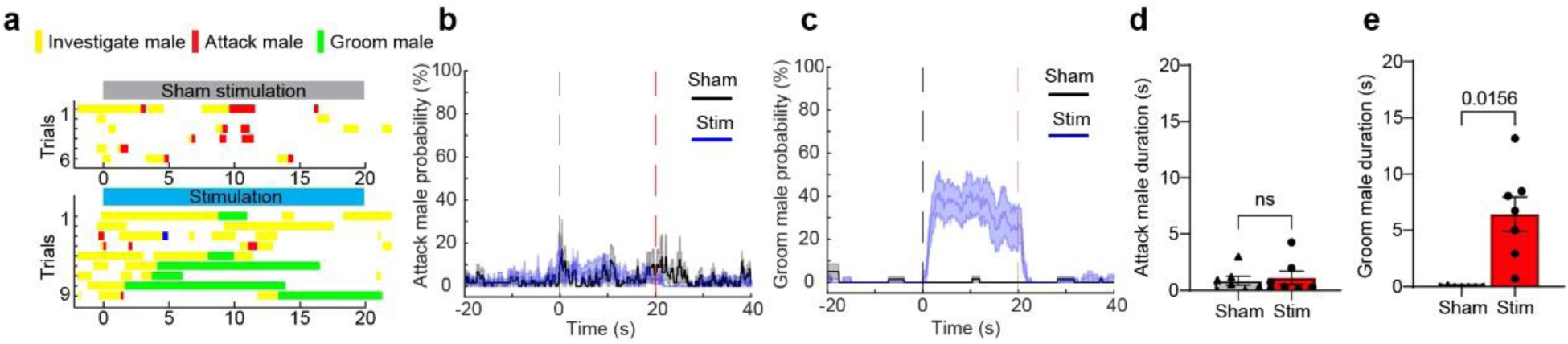
Optogenetic activation of BNSTp^Esr1^ neurons induces grooming behavior toward adult male intruders. **(a)** Raster plots showing adult male-directed behaviors during sham (top) or light (bottom) stimulation of BNSTp^Esr1^ cells in sexually naïve male mice. **(b)** PETHs of attacking adult male intruder probability aligned to sham (black) and light (blue) onset of ChR2 male mice. The dashed lines mark the trial period. (n=7 male mice. Data shown as mean ± SEM.) **(c)** PETHs of grooming adult male intruder probability aligned to sham (black) and light (blue) onset of ChR2 male mice. The dashed lines mark the trial period. (n=7 male mice. Data shown as mean ± SEM.) **(d)** The duration of attacking adult male intruders during sham and light stimulation in ChR2 male mice. (n=7 male mice. Wilcoxon test, ns: not significant; Data shown as mean ± SEM.) **(e)** The duration of grooming adult male intruders during sham and light stimulation in ChR2 male mice. (n=7 male mice. Wilcoxon test. Data shown as mean ± SEM.)

**Figure S4.**
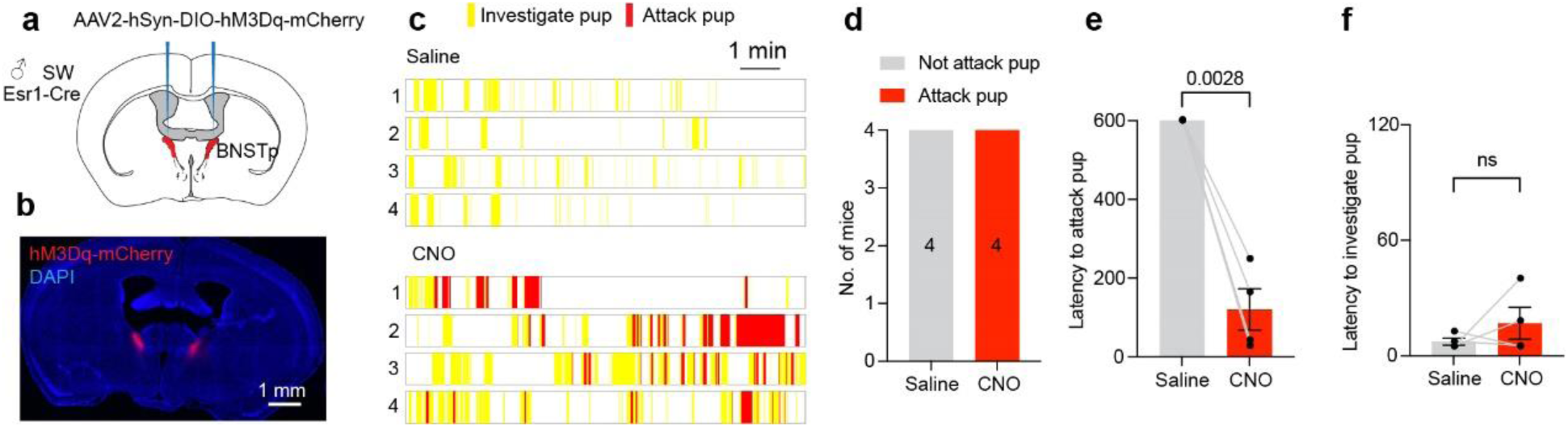
Chemogenetic activation of BNSTp^Esr1^ neurons induces male infanticide. **(a)** The strategy to chemogenetically activate BNSTp^Esr1^ cells. **(b)** A representative image showing hM4Di-mCherry (red) expression in the BNSTp. **(c)** Raster plots showing pup-directed behaviors after saline (top) and CNO (bottom) injection. **(d)** Number of males that attacked pups after saline and CNO injection. (n=4 males) **(e)** The average latency to attack pup after saline and CNO injection. The latency equals 600s if no attack occurs during the 10-minute testing period. (n=4 males. Paired t-test. Data shown as mean ± SEM.) **(f)** The average latency to investigate the pup after saline and CNO injection. (n=4 males. Paired t-test, ns: not significant. Data shown as mean ± SEM.)

